# Drivers of fish beta diversity vary by habitat and rainfall period in ephemeral aquatic habitats of the Atlantic Forest

**DOI:** 10.64898/2025.12.22.695535

**Authors:** João Henrique Alliprandini da Costa, Jansen Zuanon, Amanda Selinger, Ursulla Pereira Souza, Rafael Mendonça Duarte

## Abstract

Ephemeral aquatic habitats, such as temporary pools and roadside ditches, are characterized by cyclical flooding and drying that create highly dynamic conditions for fish community assembly. These environments can exhibit high spatial and temporal variability in community structure driven by local topography, distance to source habitats and interannual variation in rainfall, which affects connectivity dynamics. As such, they provide unique opportunities to test hypotheses about the processes shaping metacommunities, particularly through the partitioning of beta diversity into turnover and nestedness components. Here, we investigate these processes in ephemeral aquatic habitats of the Atlantic Forest, where we sampled 36 temporary pools and 36 roadside ditches throughout 2024, recording 1,545 individuals from 20 species. Fish communities’ composition did not differ significantly between habitats or hydrological periods, but beta diversity was consistently high and predominantly influenced by turnover. This suggests spatial processes such as dispersal limitation and colonization history are key drivers of community structure in those temporary aquatic habitats. We tested whether beta diversity components were associated with local environmental (temperature, dissolved oxygen, pH, volume) and spatial (distance to the nearest stream) predictors. During the dry period, nestedness was positively correlated with differences in volume for temporary pools, indicating that greater habitat availability promotes species accumulation under low connectivity, where dispersal occurs possibly only by overland displacement. Additionally, turnover was associated with differences in distance to the nearest stream in roadside ditches during the dry period, likely reflecting differential colonization by stream-associated species. During the wet period, differences in pH were correlated with turnover in roadside ditches, consistent with Mass Effect dynamics under higher connectivity conditions. Meanwhile, in temporary pools during the wet period, turnover increased with differences in distance to the nearest stream, suggesting colonization by highly tolerant and capable of dispersal species in more isolated sites. These contrasting relationships may also reflect the spatial organization of habitats, with linear ditches facilitating directional dispersal along roads, whereas spatially dispersed pools depend on more stochastic colonization. Thus, pools act as seasonal refugia for tolerant or amphibious species during low connectivity, and for stream-dwelling species during high connectivity, whereas ditches function as more persistent nodes that facilitate dispersal during wetter periods. This seasonal complementarity helps explain the observed temporal and habitat-related variation in beta diversity, highlighting that effective conservation must target the entire floodable mosaic, rather than isolated sites, to maintain this dynamic metacommunity.

## INTRODUCTION

Temporary pools are key components of freshwater landscapes in tropical regions, defined by their cyclical drying and flooding regimes, which create dynamic environments for colonization, extinction and recolonization events (Williams, 2007). Despite their small size and transient nature, they can harbor high biodiversity and play critical roles in dispersal, reproduction and life cycle for several freshwater species (Blaustein and Schwartz, 2001; DiMauro and Hunter, 2002; Espírito-Santo et al., 2013). However, temporary pools are increasingly threatened by a range of anthropogenic pressures, harboring a high number of small-sized threatened fishes (Castro and Polaz, 2020). For example, habitat loss driven by road construction, urbanization and land-use change has pushed several pool-dwelling species toward extinction (Volcan and Lanés, 2018). The disappearance of natural pools and the degradation of floodplains have also led some fishes to the occupation of roadside ditches, human-made temporary aquatic habitats built along dirt roads for water drainage, by species typically associated with temporary habitats (Costa et al., 2024, 2025). As a result, fish communities from ephemeral aquatic habitats that are naturally short-lived and structured over brief hydroperiods are now also being reshaped in artificial environmental settings that may facilitate the introduction of non-native species, together with increased exposure to pollutants and increases in natural thermal regimes (Hohausová et al., 2010; Costa et al., 2025).

Despite their ecological importance, studies on fish metacommunity in ephemeral aquatic habitats remain scarce in the Neotropical region, which harbors the richest freshwater fish fauna on the planet. Most existing community ecology studies have focused on seasonal and environmental variation in temporary pools from the Amazon and Pantanal biomes (Pazin et al., 2006; Espírito-Santo et al., 2013; Tondato et al., 2013; Espírito-Santo and Zuanon, 2017; Couto et al., 2018), with apparently no published assessments from the Atlantic Forest. Notably, the only study investigating a fish metacommunity in roadside ditches within the Neotropics was recently conducted in this biome (Costa et al., 2025).

This knowledge gap is particularly relevant because metacommunity theory provides a powerful framework for understanding how communities assemble across spatial and temporal scales. According to this theory, community assembly emerges from the interplay between environmental filtering, species interactions and dispersal processes (Leibold et al., 2004). These mechanisms are summarized in four non-mutually exclusive paradigms: (i) the Patch Dynamics paradigm emphasizes colonization–extinction dynamics among equivalent habitat patches; (ii) Species Sorting highlights the role of environmental heterogeneity and niche differentiation in determining community composition; (iii) Mass Effect describe situations in which high dispersal rates allow species to persist even in suboptimal habitats through source–sink dynamics; and (iv) Neutral Theory assumes ecological equivalence among species, with stochastic dispersal and demographic processes structuring communities. The relative importance of these paradigms varies across taxa, habitats, spatial scales and temporal contexts (Leibold et al., 2022), underscoring the need to expand metacommunity studies to different types of aquatic habitats. In this context, beta diversity, which measures spatial and/or temporal variation in communities’ composition, offers a valuable approach to investigate metacommunity structure. Its partition into turnover (species replacement) and nestedness (species loss or gain) can help identify the processes shaping assemblages (Baselga, 2010), and are fundamental for understanding how communities differ across sites and how such differences are shaped by dispersal limitation and connectivity in such spatially-disjunct aquatic habitats (Declerck et al., 2011).

Partitioning beta diversity also enables the testing of hypotheses regarding the mechanisms that structure biodiversity across local and regional scales (Legendre and De Cáceres, 2013). This is critical for conservation planning once high turnover among sites (here, a set of ephemeral aquatic habitats in a given landscape) suggests the need to protect multiple locations (in this case, ephemeral habitats), whereas high nestedness may allow prioritizing the most species-rich sites (Socolor et al., 2016). Thus, disentangling the relative contributions of turnover and nestedness to community variation in both natural and artificial ephemeral aquatic habitats is a crucial step toward addressing existing knowledge gaps in these transient and highly threatened systems. Furthermore, identifying the environmental and temporal drivers of beta diversity offers key insights into species replacement, species loss and colonization, as variation among sites within a region can be interpreted through appropriate explanatory variables (Legendre, 2014).

Ephemeral aquatic habitats of the Atlantic Forest are not only valuable systems for exploring and understanding the components and drivers of beta diversity, but also serve as replicable models for conserving metacommunities in other ecosystems. In these fragmented environments, species richness is shaped by factors such as pool volume and proximity to permanent water bodies (Pazin et al., 2006), parallel to the principles of island biogeography theory proposed by MacArthur and Wilson (1967). That is, as global landscapes become increasingly fragmented, gaining insight into beta diversity patterns in such ephemeral and extreme aquatic habitats becomes ever more critical, especially as these systems may represent the new reality for other climate-vulnerable ecosystems in the future (Comte and Olden, 2017). Therefore, in this study we investigated how fish metacommunity composition and beta diversity components (turnover and nestedness) vary across two habitat types and hydrological periods in ephemeral aquatic habitats of the Atlantic Forest. Our focus was on fish communities from temporary pools and roadside ditches sampled within a blackwater microbasin located in the coastal plains of southeastern Brazil. The region harbors a high fish endemism resulting from its geographic isolation, bounded by the Atlantic Ocean and coastal mountain ranges that restrict species dispersal between basins (Menezes et al., 2007, Giongo et al., 2023). We also evaluated how local environmental conditions that may act as ecological filters (temperature, dissolved oxygen and pH), habitat availability (using water volume as a proxy) and spatial isolation (measured as the distance to the nearest stream) influence patterns of species turnover and nestedness.

We hypothesized that, during the dry period, beta diversity components would be primarily correlated with water volume, as greater habitat availability (and diversity) inside each aquatic habitat could reduce competitive exclusion, given that connectivity is at its lowest point and dispersal is possibly only viable for fish species capable of overland displacement. In contrast, temperature, dissolved oxygen and pH were expected to have limited correlation with beta diversity components, since the species present in these ephemeral aquatic habitats were already selectively filtered for occupation of such environments. Moreover, Species Sorting is likely a mechanism acting throughout evolutionary timescales, rather than at the present ecological scale, similar to what was found in Central Amazonia, where communities were shown to be more affected by pool area and depth than by physicochemical conditions (Pazin et al., 2006). During the wet period, we predicted that none of the measured variables would be significantly correlated to beta diversity components due to enhanced connectivity among habitats, which makes environmental (physicochemical) conditions closely similar among sites. This expectation is based on findings from the Okavango Delta, where communities tend to become homogenized during high water seasons (Bokhutlo et al., 2021), consistent with a Mass Effect dynamic in which elevated immigration compensates for environmentally unsuitable conditions across the metacommunity.

## MATERIAL AND METHODS

### Study area and sampling

Sampling was conducted in the Preto River microbasin, part of the Itanhaém River basin, within the coastal Atlantic Forest of São Paulo State, Brazil (Figure 1A). This coastal plain region is characterized by sandy-clay substrates, *restinga* vegetation, and blackwater streams with high dissolved organic carbon (rich in humic substances), low pH and low dissolved oxygen (Camargo and Cancian, 2016; Table S1). From January to December 2024, 36 temporary pools (Figure 1B) and 36 roadside ditches (Figure 1C) were sampled, with three distinct sites of each habitat type surveyed monthly. A given ephemeral site, or another within the same temporarily flooded area, was only resampled in a later month if it had undergone complete drying and subsequent reflooding, or complete reconnection with streams. This criterion, as adopted by Costa et al. (2025), ensured that fish communities could recolonize naturally and reduced carryover effects from prior sampling.

**Figure 1.**
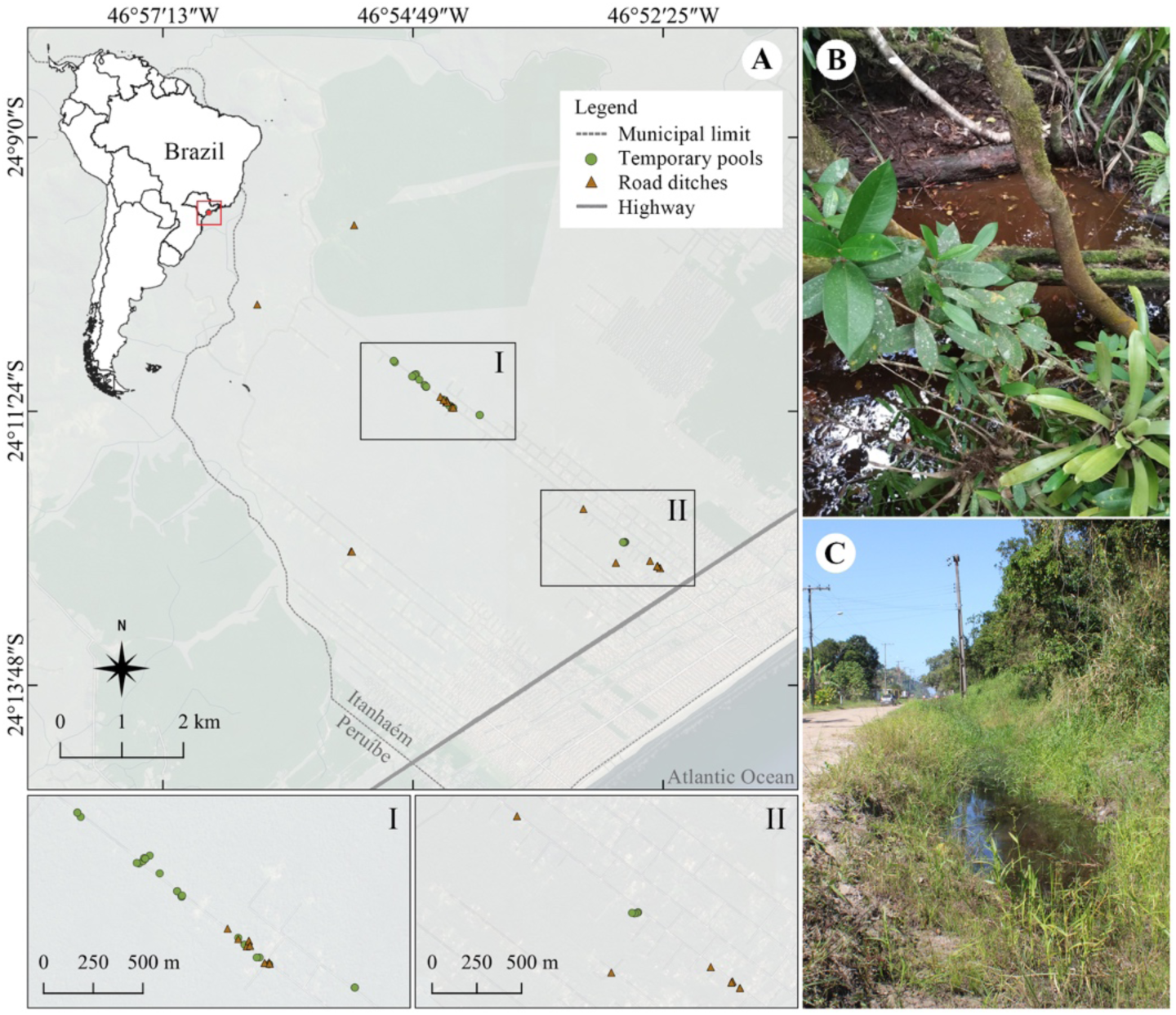
(A) Study area in the Preto River microbasin, within the Itanhaém River Basin, in the Atlantic Forest of Southeastern Brazil. Green circles represent sampled temporary pools and brown triangles represent roadside ditches. Insets I and II highlight clusters of sampled habitats located in the lowest and most flood-prone portions of the microbasin. (B) Example of a temporary pool within forested *restinga* vegetation. (C) Example of a roadside ditch.

Fish were sampled for 15 minutes at each site by two researchers using hand nets (50 × 40 cm, 1 mm mesh). Captured individuals were euthanised in eugenol (1–1.5 mL L-^1^), fixed in 10% formalin for 48 hours, and preserved in 70% ethanol. All procedures were approved by the Ethics Committee for Animal Use at São Paulo State University (UNESP – IB/CLP, protocol no. 15/2023) and authorized under SISBIO license 90241-1. Voucher specimens were deposited in the Fish Collection of UNESP in São José do Rio Preto (DZSJRP).

At each site, we measured environmental variables including maximum pool length, width, and depth (in meters) using a fiberglass measuring tape. Pool or ditch volume (m³) was estimated assuming a half-ellipsoid shape using the formula: V = 2/3 x π x a x b x c, where a is maximum length, b is maximum width, and c is maximum depth. Water temperature (°C), pH and dissolved oxygen (mg·L⁻¹) were measured in situ using a portable multiprobe (Hanna HI98194); these three variables were selected a priori for their ecological relevance in ephemeral habitats, where exposure to direct sunlight and the absence of water flow can lead to thermal and oxygen extremes, which in turn could be filtering fish assemblages. The distance from each sampling site to the nearest stream was measured using satellite imagery via Google Maps. Sampling months were grouped into two hydrological periods: the wet period (November-March) and the dry period (April-October), based on regional rainfall patterns described in Costa et al. (2025). Although we have used the term “dry” (since the rainfall difference is enough to dry most of the sampled ephemeral aquatic habitats), it is important to note that despite there being strong differences in precipitation between these periods (Costa et al., 2025), the dry period still receives rain, even in smaller quantities (see monthly water volume per aquatic habitat in Figure S1), a characteristic of the humid coastal plains of southeastern Brazil. Summary statistics (minimum, maximum, mean and standard deviation) for each environmental variable per habitat type and period are provided in Table S1, together with the number of sites sampled per habitat-period combination.

### Data analysis

To assess variation in fish assemblage composition across habitats and hydrological periods, we performed a non-metric multidimensional scaling (NMDS) based on Bray–Curtis dissimilarities calculated from species abundance data. Differences in community composition between habitat type (temporary pools and roadside ditches) and sampling period (wet and dry) were tested using Permutational Multivariate Analysis of Variance (PERMANOVA). We also tested the homogeneity of multivariate dispersion using PERMDISP to evaluate within-group variability.

Beta diversity was further quantified using the Sørensen dissimilarity index, which was partitioned into its turnover and nestedness components using the *betapart* R package (Baselga and Orme, 2012). All analyses were performed separately for each habitat-period combination using presence–absence data. To explore the drivers of beta diversity components, we calculated pairwise Euclidean distances for selected variables (temperature, dissolved oxygen, pH, volume and distance to the nearest stream) and tested their correlation with pairwise beta diversity components matrices using Mantel tests with 999 permutations. All statistical analyses were performed in R.

## RESULTS

A total of 1,545 individuals were sampled, with 758 individuals in temporary pools and 787 in roadside ditches, comprising 20 species overall. Of these, 18 species occurred in pools and 17 in ditches. The most abundant species in temporary pools were *Mimagoniates lateralis* (Nichols, 1913) (n = 281), *Atlantirivulus peruibensis* Ywamoto, Nielsen & Oliveira, 2025 (n = 157), *Hyphessobrycon boulengeri* (Eigenmann, 1907) (n = 92) and *Phalloceros reisi* Lucinda, 2008 (n = 79). In roadside ditches, the dominant species were *A. peruibensis* (n = 212), *H. boulengeri* (n = 131), *P. reisi* (n = 104), *Poecilia reticulata* Peters, 1859 (n = 77) and *Hyphessobrycon bifasciatus* Ellis, 1911 (n = 72) (Table 1). We identified two confirmed non-native species, *P. reticulata* was exclusive to roadside ditches, while *Nannostomus beckfordi* Günther, 1872 occurred in both aquatic habitats, but with greater abundance in ditches. A single specimen of a possible third non-native species, *Hoplosternum littorale* (Hancock, 1828), also occurred in a roadside ditch; however, it’s origin as native or non-native to the southeastern Brazil basins is not fully understood yet. In addition, we recorded three endangered species: *Leptopanchax itanhaensis* (Costa, 2008), classified as Critically Endangered, and more abundant on roadside ditches than in temporary pools (Table 1); *Rachoviscus crassiceps* Myers, 1926 and *M. lateralis*, both classified as Vulnerable, with *R. crassiceps* more abundant in ditches and *M. lateralis* more abundant in pools. Fish assemblages’ composition did not differ significantly between habitats or across sampling periods (PERMANOVA: F_3,68_ = 1.333, R² = 0.056, p = 0.085; PERMDISP: F_3,68_ = 0.292, p = 0.831; Figure 2).

**Figure 2.**
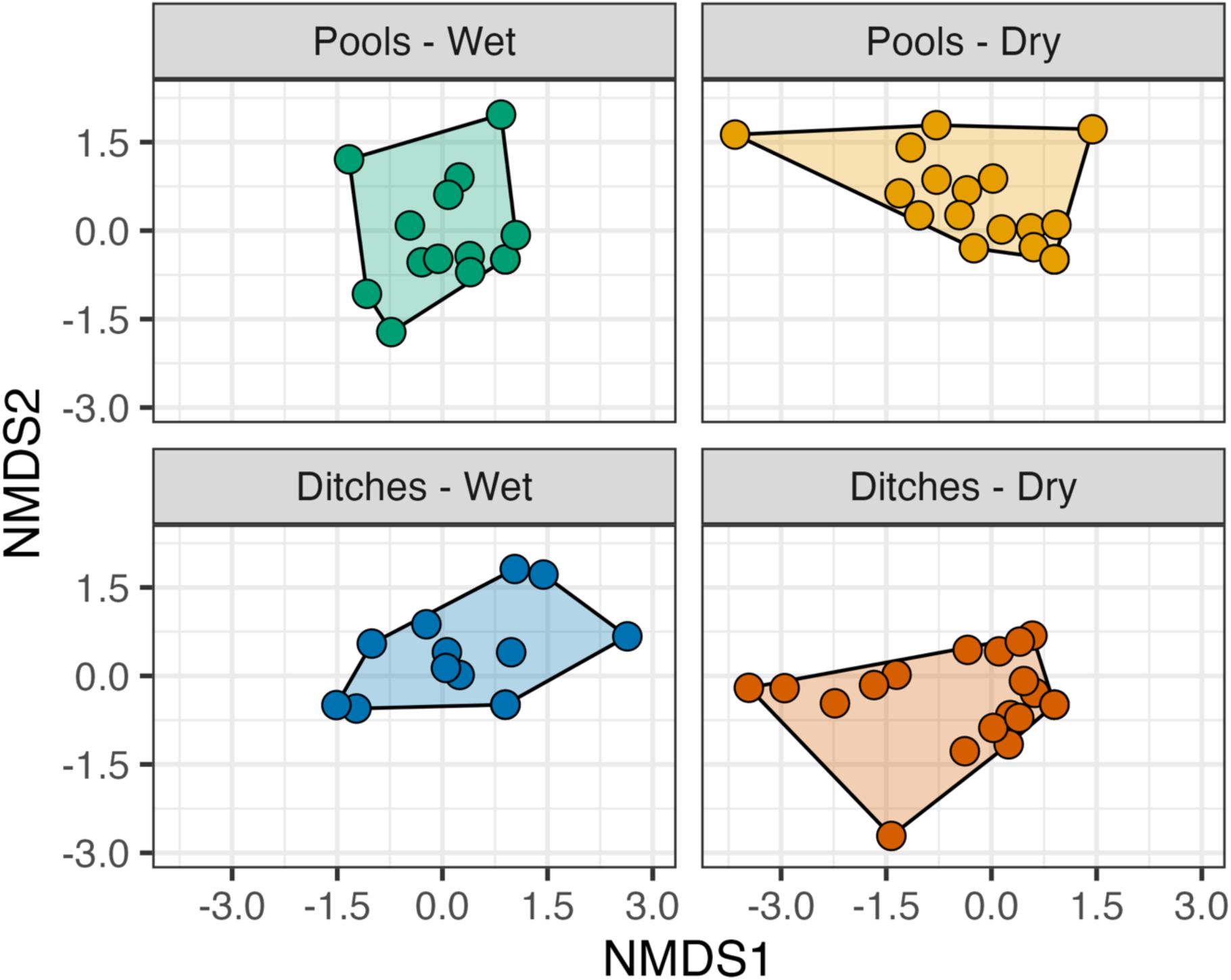
Non-metric multidimensional scaling (NMDS) plots based on Bray–Curtis dissimilarities of fish species abundance data, showing assemblage structure across habitat type and sampling period in ephemeral aquatic habitats of the Preto River microbasin, in the Atlantic Forest of Southeastern Brazil. Convex hulls represent the multivariate space occupied by each group.

**Table 1.** Fish species recorded in temporary pools (TP) and roadside ditches (RD) of the Preto River microbasin, in the Atlantic Forest of Southeastern Brazil. Values indicate total abundance per habitat type across all samples. Voucher specimens are deposited in the DZSJRP collection of UNESP. * non-native species.

| Orders and families | Species | TP | RD | Voucher<br>DZSJRP |
| --- | --- | --- | --- | --- |
| CHARACIFORMES |  |  |  |  |
| Stevardiidae |  |  |  |  |
|  | *Mimagoniates lateralis* (Nichols, 1913) | 281 | 62 | 25059 |
| Acestrorhamphidae |  |  |  |  |
|  | *Hyphessobrycon boulengeri* (Eigenmann, 1907) | 92 | 131 | 25058 |
|  | *Hyphessobrycon bifasciatus* Ellis, 1911 | 1 | 72 | 25070 |
|  | *Rachoviscus crassiceps* Myers, 1926 | 7 | 9 | 23340 |
|  | *Hollandichthys multifasciatus* (Eigenmann & Norris, 1900) | 41 | 1 | 25062 |
| Crenuchidae |  |  |  |  |
|  | *Characidium lanei* Travassos, 1967 | 4 | 0 | 25061 |
| Erythrinidae |  |  |  |  |
|  | *Hoplias malabaricus* (Bloch, 1794) | 2 | 8 | 25065 |
| Lebiasinidae |  |  |  |  |
|  | *Nannostomus beckfordi* Günther, 1872* | 3 | 35 | 25069 |
| CICHLIFORMES |  |  |  |  |
| Cichlidae |  |  |  |  |
|  | *Geophagus iporangensis* Haseman, 1911 | 12 | 4 | 25066 |
| CYPRINODONTIFORMES |  |  |  |  |
| Poeciliidae |  |  |  |  |
|  | *Phalloceros reisi* Lucinda, 2008 | 79 | 104 | 25064 |
|  | *Poecilia reticulata* Peters, 1859* | 0 | 77 | 25068 |
| Rivulidae |  |  |  |  |
|  | *Atlantirivulus peruibensis* Ywamoto, Nielsen & Oliveira, 2025 | 157 | 212 | 25056 |
|  | *Leptopanchax itanhaensis* (Costa, 2008) | 1 | 13 | 24845; 24846 |
| GYMNOTIFORMES |  |  |  |  |
| Gymnotidae |  |  |  |  |
|  | *Gymnotus pantherinus* (Steindachner, 1908) | 25 | 0 | 25067 |
| SILURIFORMES |  |  |  |  |
| Callichthyidae |  |  |  |  |
|  | *Callichthys callichthys* (Linnaeus, 1758) | 9 | 49 | 25057 |
|  | *Hoplosternum littorale* (Hancock, 1828)* | 0 | 1 | 25173 |
|  | *Scleromystax macropterus* (Regan, 1913) | 35 | 1 | 25060 |
| Loricariidae |  |  |  |  |
|  | *Pseudotothyris obtusa* (Miranda Ribeiro, 1911) | 6 | 6 | 25073 |
| Heptapteridae |  |  |  |  |
|  | *Acentronichthys leptos* Eigenmann & Eigenmann, 1889 | 2 | 0 | 25071 |
| SYNBRANCHIFORMES |  |  |  |  |
| Synbranchidae |  |  |  |  |
|  | *Synbranchus marmoratus* Bloch, 1795 | 1 | 2 | 25072 |

Beta diversity was high in all groups (β_Sørensen_ ≥ 0.86), with species turnover as the dominant component (β_turnover_ ≥ 0.71) and a smaller contribution from nestedness (β_nestedness_ ≤ 0.18). This pattern was consistent across habitats and hydrological periods (Figure 3). When comparing sites pairwise within each habitat-period combination, we found that differences in water volume were significantly associated with nestedness for temporary pools during the dry period (Mantel r = 0.44, p = 0.001; Figure 4A). Additionally, differences in distance to the nearest stream were positively correlated with turnover in temporary pools during the wet period (r = 0.42, p = 0.01; Figure 4B) and in roadside ditches during the dry period (r = 0.26, p = 0.017; Figure 4D). A significant correlation was also found for differences in pH with species turnover for roadside ditches during the wet period (r = 0.37, p = 0.007, Figure 4C). No significant associations were detected between pairwise beta diversity components with differences in temperature and dissolved oxygen (Table 2).

**Figure 3.**
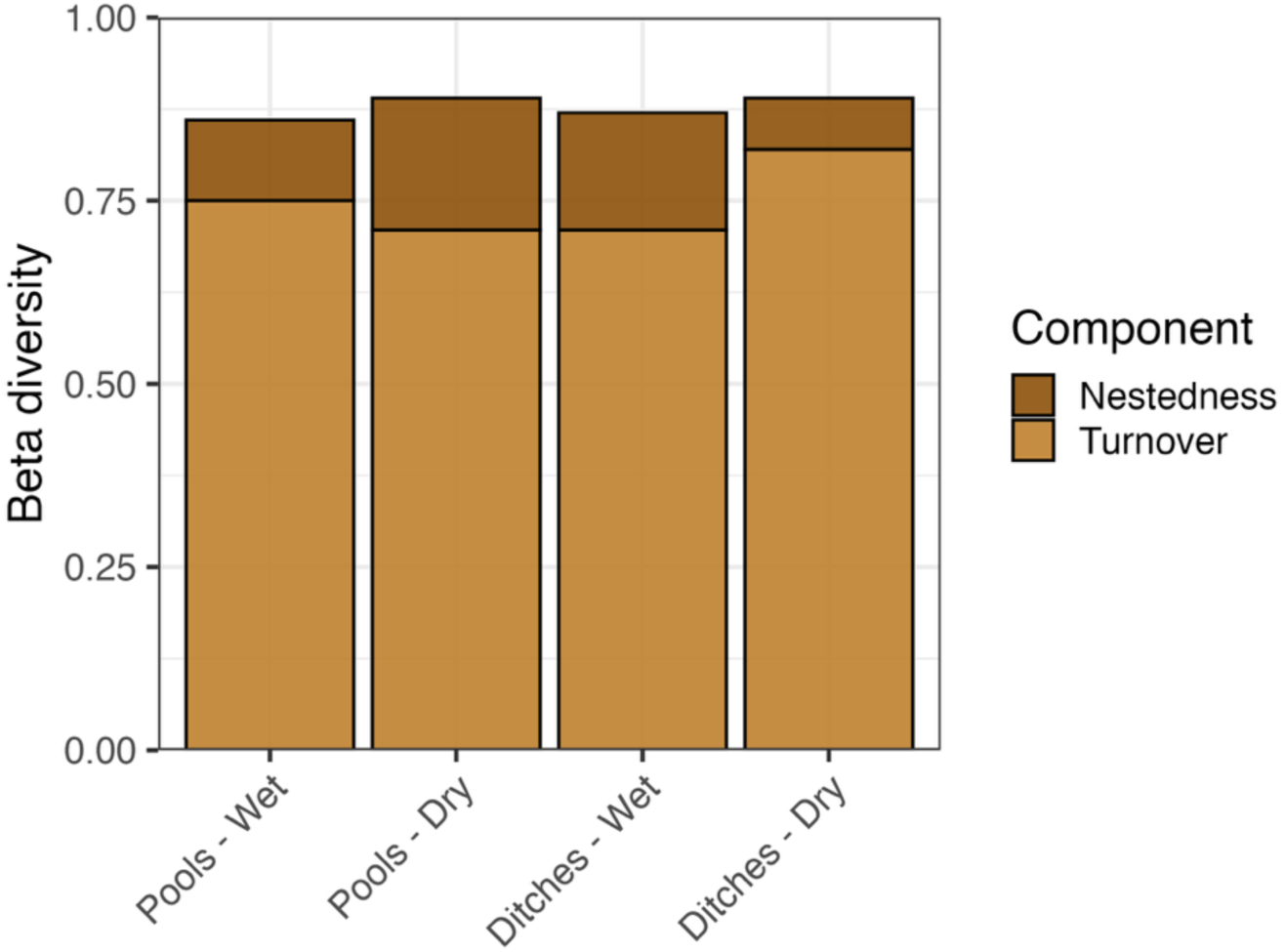
Partitioning of beta diversity (Sørensen index) into turnover and nestedness components for fish assemblages in ephemeral aquatic habitats of the Preto River microbasin, in the Atlantic Forest of Southeastern Brazil.

**Figure 4.**
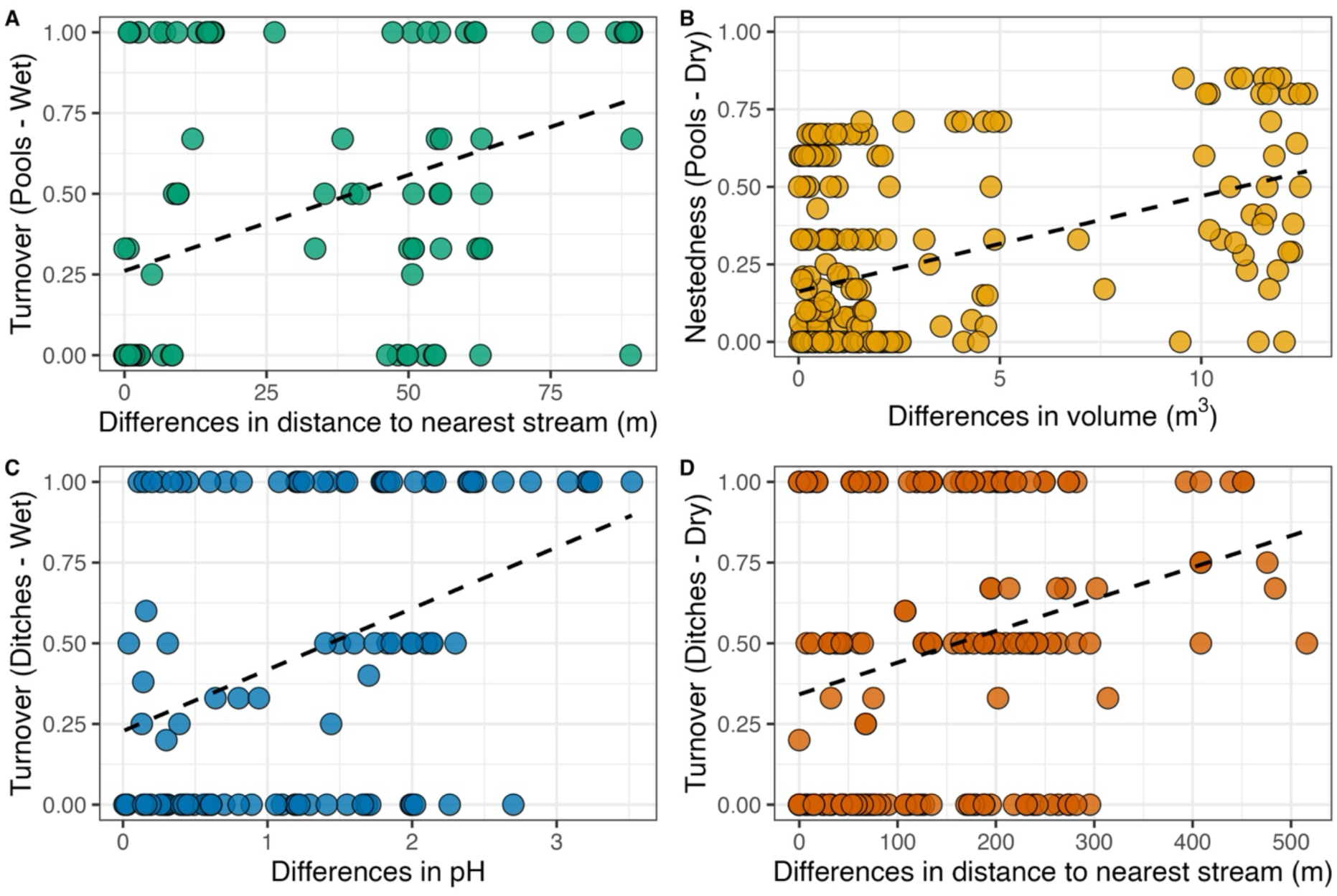
Significant relationships between predictors distances and pairwise beta diversity components in ephemeral aquatic habitats of the Preto River microbasin, in the Atlantic Forest of Southeastern Brazil. (A) Positive correlation between difference in distance to nearest stream and turnover in temporary pools during the wet period. (B) Positive correlation between difference in volume and nestedness in temporary pools during the dry period. (C) Positive correlation between difference in pH and turnover in roadside ditches during the wet period. (D) Positive correlation between difference in distance to nearest stream and turnover in roadside ditches during the dry period. Dashed lines represent linear trends for visualization.

**Table 2.**
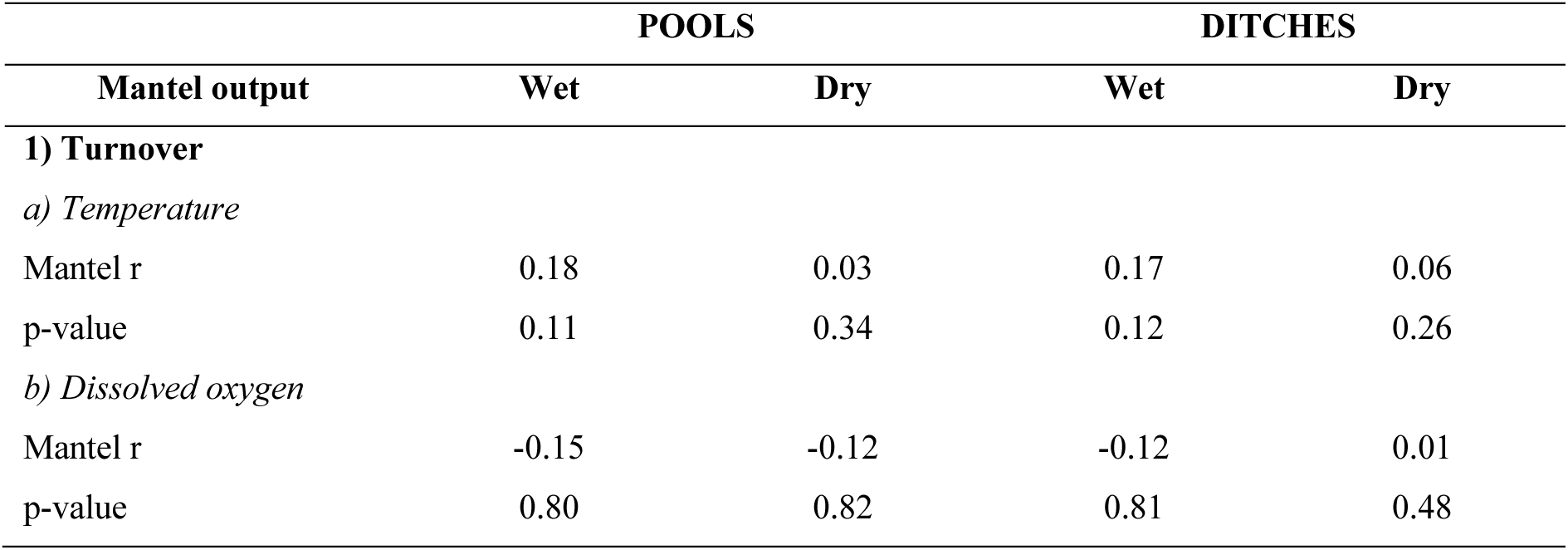
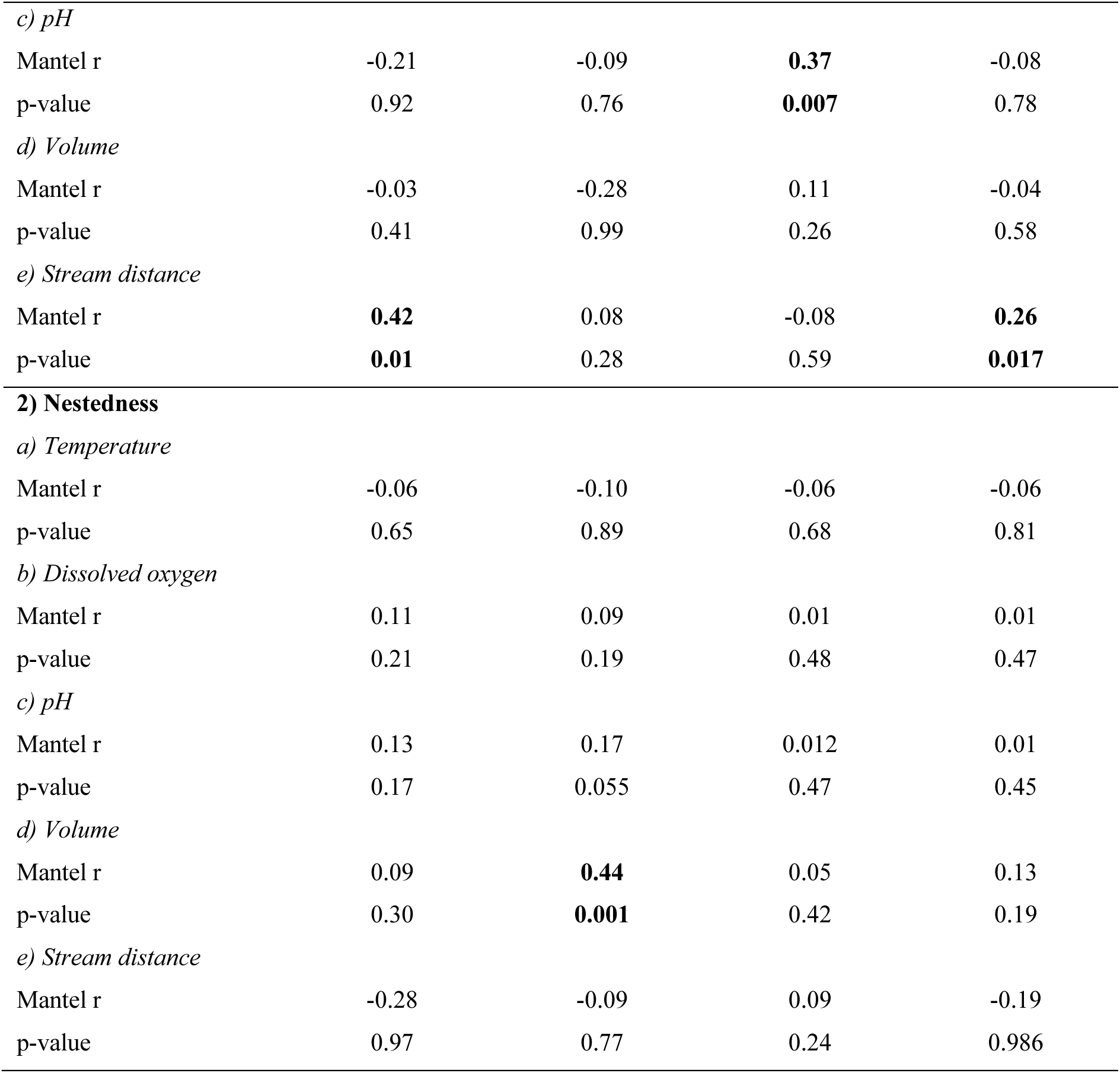
Mantel test results showing correlations (r) and associated p-values between pairwise beta diversity components (turnover and nestedness) and predictors dissimilarities (temperature, dissolved oxygen, pH, volume and distance to the nearest stream) across habitat and sampling period of ephemeral aquatic habitats from the Preto River microbasin, in the Atlantic Forest of Southeastern Brazil. Significant results (p < 0.05) are shown in bold.

By analyzing these significant relationships according to the distribution of each species (Figure 5), it is possible to see that the relationship between difference in distance to the nearest stream and turnover for temporary pools during the wet period was mostly driven by the appearance of *Callichthys callichthys* (Linnaeus, 1758), *Gymnotus pantherinus* (Steindachner, 1908), *Hoplias malabaricus* (Bloch, 1794) and *R. crassiceps* in more distant pools, while some species were more abundant in pools very close to streams, such as *Acentronichthys leptos* Eigenmann & Eigenmann, 1889 and *Hollandichthys multifasciatus* (Eigenmann & Norris, 1900) (Figure 5A). Meanwhile, the relationship between difference in volume and nestedness for temporary pools during the dry period appears to reflect the addition of species in pools with greater volume, such as *H. malabaricus*, *H. bifasciatus* and the non-native *N. beckfordi*, and the disappearance of some species, such as *Synbranchus marmoratus* Bloch, 1795, *C. callichthys* and *A. leptos*, as pool volume increased (Figure 5B). The relationship between pH differences and turnover in roadside ditches during the wet period appears to reflect the (possibly overland) displacement of *A. peruibensis* in ditches with higher pH (that is, more filled by rain water than stream-flooded ditches), along with stream-dwelling species, such as *H. boulengeri*, *M. lateralis* and *Scleromystax macropterus* (Regan, 1913), that became more abundant in ditches with higher pH, while *R. crassiceps*, *H. littorale*, *H. multifasciatus* and *S. marmoratus* were found only in acidic pH (Figure 5C). For roadside ditches during the dry period, the relationship found between the difference in distance to the nearest stream and turnover appears to be related to the replacement of stream-dwelling species by non-native species, *P. reticulata* and *N. beckfordi*, along with the native *P. reisi*, the farther from the stream (Figure 5D).

**Figure 5.**
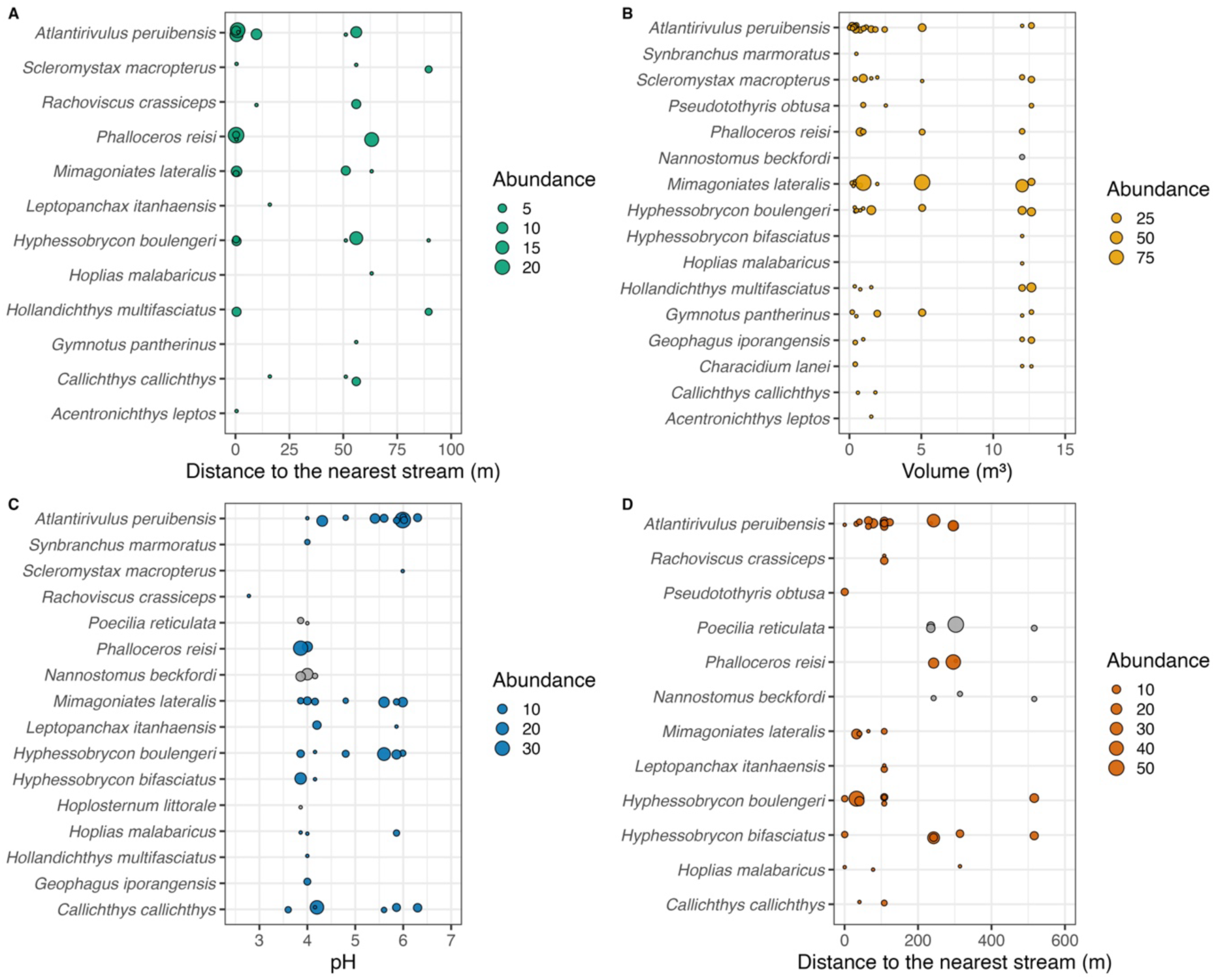
Species abundance according to environmental and spatial isolation gradients that had significant correlations with beta diversity components. (A) Species abundance along the distance to the nearest stream (m) in temporary pools during the wet period. (B) Species abundance along the water volume (m^3^) in temporary pools during the dry period. (C) Species abundance along the pH in roadside ditches during the wet period. (D) Species abundance along the distance to the nearest stream (m) in roadside ditches during the dry period. Non-native species are colored gray.

## DISCUSSION

Our study represents the first effort to investigate beta diversity components of a fish metacommunity in temporary pools and roadside ditches of the Atlantic Forest, an important step toward understanding and conserving the fauna of these ephemeral aquatic habitats in one of the last conserved remnants of the biome. It also provides the first report of fish communities in temporary pools from this biome, complementing recent findings on roadside ditches (Costa et al., 2025). We found a similar total number of individuals and species in both habitat types throughout the year, with *A. peruibensis* emerging as the second most abundant species in pools and the most abundant in ditches. This pattern aligns with the species’ highly amphibious lifestyle, including terrestrial locomotion via leaping, which facilitates both escape from predators and colonization of new ephemeral habitats, as evidenced in other Rivulidae species (Espírito-Santo et al., 2019). However, in the pools, *A. peruibensis* was outnumbered by *M. lateralis*, a typical stream-dwelling tetra endemic from the region (Moraes et al., 2020; Selinger et al., 2025), probably by the fact that *M. lateralis* occurs mainly in large schools in floodplain pools, with up to 90 individuals recorded at a single site.

A key finding was the absence of significant differences in species composition between habitat types or hydrological periods. This suggests that fish communities are similar across these ephemeral habitats, regardless of their origin (natural or man-made) or seasonal variation. This pattern is consistent with findings from ephemeral aquatic systems in subtropical Africa, where hydrological regimes act as major drivers of community structure and increased connectivity during wet periods promotes assemblage homogenization (Bokhutlo et al., 2021). In addition, in southeastern Brazil, even the dry period includes unregular rainfall events, and not all pools or ditches dry synchronously and completely, as evidenced by Figure S1. As a result, some habitats retain both pool-adapted species capable of tolerating harsh conditions, and trapped stream-associated species such as *M. lateralis*, which likely contributes to the observed compositional similarity. Nonetheless, certain species were exclusive to each habitat type. *Characidium lanei* Travassos, 1967, *G. pantherinus* and *A. leptos* were found only in temporary pools, while *H. littorale* and *P. reticulata* occurred exclusively in roadside ditches.

The presence of *P. reticulata*, a non-native species introduced for mosquito control and known for its broad environmental tolerance (Silva et al., 2023), together with *N. beckfordi*, a species native to the Amazon basin that has been widely exported as an ornamental fish and introduced in various regions (Magalhães et al., 2019), with both species occurring mostly in ditches, is of particular concern. Although *P. reticulata* has not yet been observed in temporary pools, continued monitoring is essential to assess its potential spread and long-term impact on these vulnerable ecosystems, and it’s probably just a matter of detection, since enhanced connectivity during the wet period has the potential to disperse these non-native species between aquatic habitats, even though roadside ditches probably are a first introduction pathway. Another noteworthy species is *H. littorale*. Although only one specimen was found in a roadside ditch, its occurrence as native or non-native in southeastern Brazil needs to be further investigated in future studies, since records for the species appear to exist only in recent decades for the region, with a history of only one deposited specimen from the upper Paraná River in the 90s (Reis, 1997), and it is already reported as non-native in other Brazilian river basins (Moreira et al., 2023). For streams in the Itanhaém River basin, only six individuals have been reported in the past (Ferreira and Petere, 2009), which could reflect a recent dispersal from the upper Paraná River to coastal basins. Furthermore, local residents from the Preto River microbasin consume the species by collecting it in streams (JHAC, personal observation). That is, not only does the species, if non-native, have the potential to compete with *C. callichthys*, a native species with a similar niche space, altering the dynamics of these ephemeral aquatic habitats, but it may also already have cultural value for local communities. However, future research should investigate the occurrence of *H. littorale* in southeastern Brazil, with the aim of clarifying its native or non-native status.

Moreover, our findings demonstrate that fish beta diversity in ephemeral aquatic habitats of the Atlantic Forest is consistently high and predominantly driven by species turnover, regardless of habitat type or hydrological period. This pattern indicates that species replacement is the primary process structuring fish assemblages across space and time in these systems. Turnover is often associated with environmental filtering and biotic interactions (Legendre and De Cáceres, 2013), yet previous research (Pazin et al., 2006; Bokhutlo et al., 2021; Costa et al., 2025), along with our results (particularly the lack of significant correlations with temperature and dissolved oxygen), suggests that limnological variables currently do not act as strong filters in these environments (Leibold and Chase, 2017). Since the species present in these ephemeral aquatic habitats were already (historically) filtered for occupation of such environments with high natural oscillations in temperature and dissolved oxygen, Species Sorting may instead operate at broader evolutionary scales, potentially influencing beta diversity over longer temporal dimensions (Jovanovska et al., 2022).

The consistently high turnover observed, even within a system of relatively low local richness, further supports this interpretation. Although the Itanhaém River Basin harbors around 64 fish species in total, considering streams (Ferreira and Petrere, 2009), only 20 species were recorded in the temporary pools and roadside ditches surveyed in the Preto River microbasin, some not reported in Ferreira and Petrere (2009), such as *L. itanhaensis*, *R. crassiceps* and *P. reticulata*. This contrast suggests that the fish fauna occupying ephemeral aquatic habitats represents a specialized subset of the regional metacommunity, adapted to tolerate harsh and fluctuating environmental conditions. Such restricted occurrences, coupled with small habitat size and low connectivity, likely intensifies competitive interactions, also contributing to the high turnover observed.

This pattern is consistent with our hypothesis that physicochemical variables would have limited influence on beta diversity components, with the sole exception of roadside ditches during the wet period, where differences in pH were positively correlated with turnover. Rather than reflecting Species Sorting through environmental filtering, this relationship seems to indicate Mass Effect dynamics, in which intense rainfall and increased hydrological connectivity allow species with good dispersal capacities to temporarily occupy suboptimal habitats with less acidic pH levels, atypical for the basin. These conditions do not seem to exclude acid-tolerant taxa, what would be expected in an environmental filter, as species such as *L. itanhaensis*, *M. lateralis* and *S. macropterus*, all typically found in highly acidic waters, were also recorded in these sites. Likewise, *H. boulengeri* and *M. lateralis*, which are extremely abundant in the microbasin (Ferreira et al., 2014), likely colonize these habitats through increased dispersal during wet periods. Additionally, this turnover appears to have been highly influenced by the occurrence of *A. peruibensis* and *C. callichthys*, two species highly tolerant to emersion, capable of terrestrial locomotion (Le Bail et al., 2000) and able to colonize rain-fed habitats independently of stream connectivity.

We also hypothesized that during the wet period no variable would be correlated with beta diversity components, due to the homogenization of the metacommunity through enhanced connectivity. However, in addition to what was evidenced by difference in pH, an exception was also observed for temporary pools: turnover was significantly correlated with differences in distance to the nearest stream. This pattern may reflect differential colonization dynamics, where pools closer to streams are more readily accessed by stream-dwelling species, while more isolated pools may be occupied by species with unique adaptations, such as *C. callichthys*, a catfish that can disperse across terrestrial environments and breathe air via a vascularized intestine (Le Bail et al., 2000), enabling it to colonize and persist in distant pools. Thus, the observed turnover reflects spatial variation in colonization sources driven by proximity to permanent water bodies and by species-specific dispersal abilities across the floodplain. However, it is important to emphasize that this pattern in the wet period was found for the year we measured, and that it may depend on inter-annual climatic variations, mainly in precipitation (that influences directly flood capacity), which should be investigated in long-term studies to verify consistency in metacommunity structuring. A similar relationship with stream distance was also found in roadside ditches during the dry period, but the underlying mechanism might differ. Once ditches are spatially linear and aligned with dirt roads, they may have been colonized historically in sequence from the sites closest to streams, functioning as stepping stones that facilitate dispersal and gradual occupation along their length. This directional connectivity could maintain turnover even under reduced hydrological connection, contrasting with the more irregular and spatially patchy distribution of temporary pools, where colonization likely depends on local stochastic events rather than sequential expansion. Moreover, more isolated ditches may favor non-native species such as *P. reticulata* and *N. beckfordi*, since introductions often occur in the most distant or anthropized ditches, that are closer to urban areas, suggesting a reversed stepping stone dynamic, where non-native species disperse toward more connected sites, contrasting with the colonization flow from ditches spatially close to streams. This process likely contributes to the observed compositional differentiation along the distance to the nearest stream gradient.

In contrast, during the dry period competitive exclusion may play a more immediate role. In resource-limited and spatially constrained habitats, distinct species among sites may reduce niche overlap and alleviate interspecific competition, as evidenced by temporary pools, supporting our hypothesis on habitat availability reducing competition, where distance in volume was significantly associated with nestedness. The absence of this pattern in roadside ditches during the dry period may be due to their inherently greater water retention, which buffers against extreme drying even during the dry months, thereby reducing variation in habitat availability across sites. In the temporary pools, the nestedness-volume significant relationship appears to reflect the addition of certain stream-dwelling species in pools with increased volume, such as *H. malabaricus* and *H. bifasciatus*, as well as the non-native *N. beckfordi*, but with the disappearance of species such as *C. callichthys*, *S. marmoratus* and *A. leptos*. This pattern is possible because these added Characiformes likely have higher niche overlap with phylogenetically close species than the species removed with increased volume, which are functionally unique to the region, with no congeners (Ferreira et al., 2014; Giongo et al., 2023). Therefore, increased habitat availability would allow niche partitioning between these similar species, facilitating species accumulation and promoting coexistence (Taylor, 1997; Spencer et al., 1999). Future studies should investigate how microhabitat heterogeneity and trait similarity vary across habitat size gradients to better understand niche partitioning in such ephemeral systems. Smaller ephemeral aquatic habitats likely support only highly amphibious or tolerant to hypoxic conditions species, such as *A. peruibensis* (Costa et al., 2025), while larger sites may host more functionally diverse assemblages, but also with greater functional redundancy, which still needs to be further investigated.

In summary, our study provides the first insights into beta diversity components in an ephemeral aquatic habitats fish metacommunity of the Atlantic Forest, revealing that species turnover predominates across both natural and man-made environments, regardless of hydrological season. This consistent pattern underscores the importance of spatial processes, particularly dispersal limitation (or the opposite, enhanced connectivity) and habitat availability, in shaping community structure. In the meantime, environmental filtering via physicochemical variables appears weak, but also suggesting that Mass Effect and local habitat features (such as habitat availability and spatial isolation), especially during low-connectivity periods, play a role in promoting coexistence or driving species replacement and nestedness.

### Implications for conservation

The consistently high turnover observed across ephemeral habitats readily suggests that conservation strategies should focus on protecting multiple sites rather than prioritizing only species-rich locations (Socolar et al., 2016). However, applying this principle to temporary pools and roadside ditches is challenging due to their inherently dynamic nature. Pools and water-filled ditches can form, dry, and reappear in different locations, with distant shapes and volumes, and often with distinct species compositions depending on connectivity patterns and stochastic colonization events, as shown for roadside ditches by our present data and by Costa et al. (2025). This spatial and temporal variability implies that protecting isolated sites is unfeasible. Instead, conservation efforts should target the entire floodable mosaic within a basin or microbasin, encompassing the naturally floodable forest areas where these habitats form and reconfigure over time. Such a systemic approach is particularly urgent in the Preto River microbasin, since Brazilian Law 12.651/2012 (Art. 61-A) (the so-called Brazilian Forest Code) designates Permanent Preservation Areas only along narrow strips, 20 to 30 meters on each side for small streams (<10 m of width), intended to protect water resources and biodiversity. This legal framework, even if strictly enforced, would fail to safeguard many pools that arise beyond these strips in floodable forests, as well as virtually all fish assemblages inhabiting roadside ditches along dirt roads. This gap reveals the inadequacy of current regulations to encompass the ecological reality of ephemeral aquatic habitats both in pristine and anthropogenically altered landscapes.

Given these limitations, a more ambitious conservation strategy is needed, one that explicitly recognizes ephemeral habitats as integral components of coastal freshwater systems. This could involve expanding the width of legally protected floodplain vegetation to include the full inundation zone of microbasins, effectively buffering both natural and man-made ephemeral sites. In the case of roadside ditches, management actions should focus on monitoring and preventing the spread of non-native species, while also avoiding road paving or ditch channelization, which would disrupt their role as temporary refugia and recolonization sources during extreme periods. Although this study focuses on a single microbasin, extending similar assessments to other coastal basins is essential. Given the high turnover observed within and likely between basins, a network of smaller protected areas across multiple basins may be more effective than concentrating protection within a single large area, as it would better capture the spatial complementarity of fish metacommunities in ephemeral aquatic habitats. Such an approach not only aligns with the ecological patterns observed but also challenges the current scope of the Forest Code, highlighting the need to revise or reinterpret existing legal instruments to ensure the conservation of these dynamic and overlooked environments.

## DATA AVAILABILITY STATEMENT

The step-by-step analysis is available in R code, along with the raw data, in a Git Hub repository (https://github.com/JH-All/ephemeral_aquatic_habitats).

## ACKNOWLEDGMENTS

We would like to thank everyone on the team who helped during the fieldwork, and Antônio Carlos (Tuca) who welcomed us into his home monthly to develop the fieldwork.

