## Supplementary figure and table for "Drivers of fish beta diversity vary by habitat and rainfall period in ephemeral aquatic habitats of the Atlantic Forest"

**SUPPLEMENTARY MATERIAL**

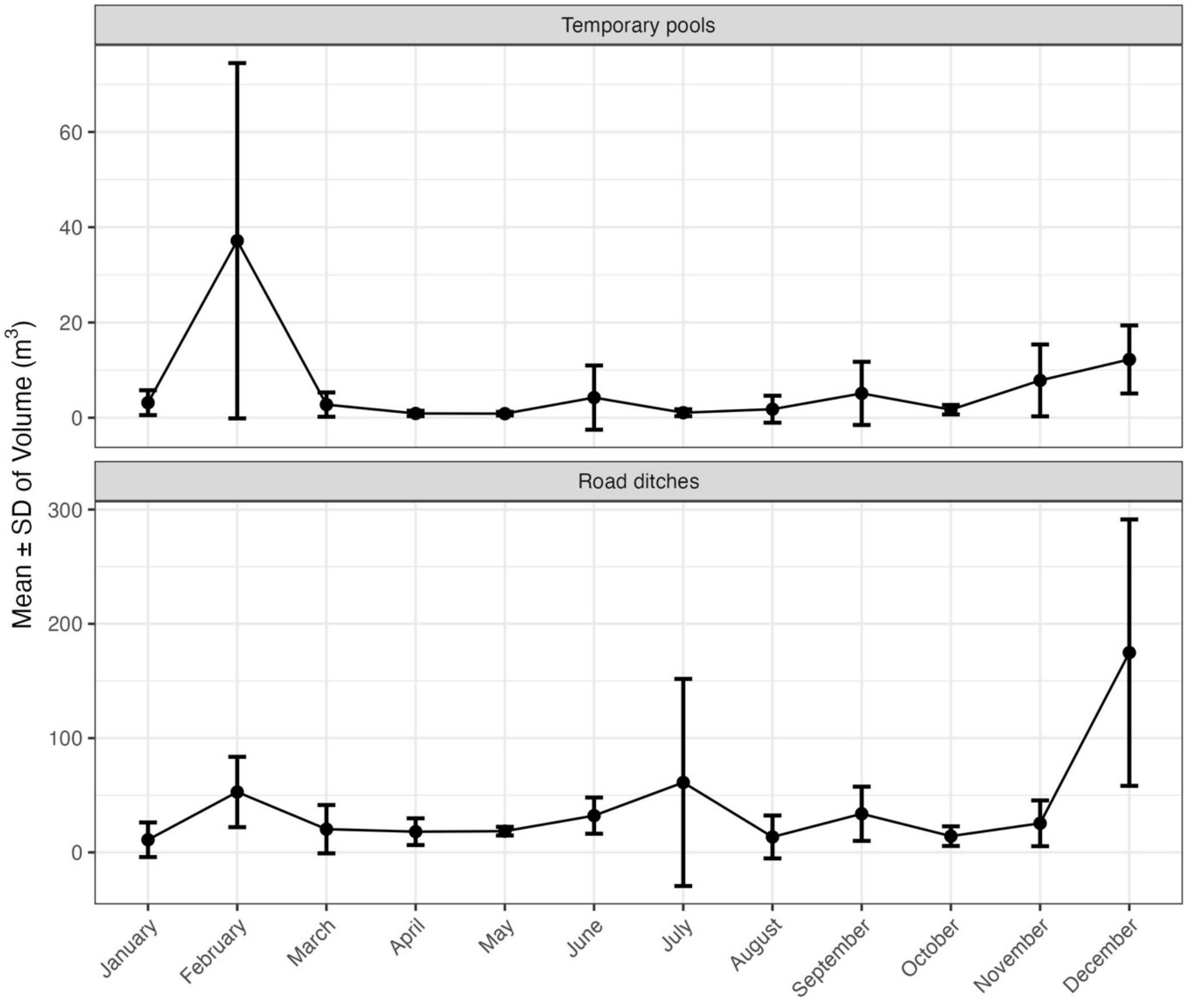


**Figure S1.** Monthly variation in mean ± standard deviation (SD) water volume ($m^3$) of temporary pools and roadside ditches in the Preto River microbasin, in the Atlantic Forest of Southeastern Brazil, during 2024.

**Table S1.** Summary statistics for environmental variables in each habitat-period category. Shown are the number of sites sampled, and the minimum, maximum, mean and standard deviation for temperature, dissolved oxygen, pH, volume and distance to the nearest stream. SD = Standard Deviation

| Variable | Statistic | Pools - Wet | Pools - Dry | Ditches - Wet | Ditches - Dry |
|---|---|---|---|---|---|
| **Number of sites** | - | 15 | 21 | 15 | 21 |
| **Temperature (°C)** | Minimum | 20.5 | 15.7 | 20.9 | 16.1 |
| | Maximum | 28.3 | 24.2 | 33 | 27.1 |
| | Mean | 25 | 20.2 | 26.8 | 21.5 |
| | SD | 2.3 | 2.68 | 3.11 | 3 |
| **Dissolved oxygen ($mg.L^{-1}$)** | Minimum | 0.0 | 0.0 | 0.0 | 0.0 |
| | Maximum | 5.15 | 2.89 | 5.2 | 2.62 |
| | Mean | 1.51 | 0.59 | 1.74 | 0.34 |
| | SD | 1.45 | 1.02 | 1.75 | 0.78 |
| **pH** | Minimum | 3.1 | 3.02 | 2.78 | 3.33 |
| | Maximum | 6.04 | 5.89 | 6.3 | 6.49 |
| | Mean | 4.39 | 3.89 | 4.86 | 5.2 |
| | SD | 0.89 | 0.91 | 1.09 | 1.03 |
| **Volume ($m^3$)** | Minimum | 0.05 | 0.03 | 1.49 | 0.63 |
| | Maximum | 77.4 | 12.7 | 293.4 | 165.37 |
| | Mean | 12.63 | 2.23 | 56.9 | 27.4 |
| | SD | 19.8 | 3.55 | 78.5 | 34.9 |
| **Distance to nearest stream (m)** | Minimum | 0.27 | 0.2 | 0.3 | 0.3 |
| | Maximum | 89.6 | 90 | 851 | 516 |
| | Mean | 19.6 | 18.2 | 141 | 179 |
| | SD | 29.7 | 29.1 | 216 | 128 |
